# A trafficome-wide RNAi screen reveals deployment of early and late secretory host proteins and the entire late endo-/lysosomal vesicle fusion machinery by intracellular *Salmonella*

**DOI:** 10.1101/848549

**Authors:** Alexander Kehl, Vera Göser, Tatjana Reuter, Viktoria Liss, Maximilian Franke, Christopher John, Christian P. Richter, Jörg Deiwick, Michael Hensel

## Abstract

The intracellular lifestyle of *Salmonella enterica* is characterized by the formation of a replication-permissive membrane-bound niche, the *Salmonella*-containing vacuole (SCV). A further consequence of the massive remodeling of the host cell endosomal system, intracellular *Salmonella* establish a unique network of various *Salmonella*-induced tubules (SIT). The bacterial repertoire of effector proteins required for the establishment for one type of these SIT, the *Salmonella*-induced filaments (SIF), is rather well-defined. However, the corresponding host cell proteins are still poorly understood. To identify host factors required for the formation of SCV and SIF, we performed a sub-genomic RNAi screen. The analyses comprised high-resolution live cell imaging to score effects on SIF induction, dynamics and morphology. The hits of our functional RNAi screen comprise: i) The late endo-/lysosomal SNARE (soluble *N*-ethylmaleimide-sensitive factor attachment protein receptor) complex, consisting of STX7, STX8, VTI1B, and VAMP7 or VAMP8, this is, in conjunction with RAB7 and the homotypic fusion and protein sorting (HOPS) tethering complex, a complete vesicle fusion machinery. ii) Novel interactions with the early secretory GTPases RAB1A and RAB1B, possibly providing a link to coat protein complex I (COPI) vesicles and reinforcing recently identified ties to the endoplasmic reticulum. iii) New connections to the late secretory pathway and/or the recycling endosome via the GTPases RAB3A, RAB8A, and RAB8B and the SNAREs VAMP2, VAMP3, and VAMP4. iv) An unprecedented involvement of clathrin-coated structures. The resulting set of hits allowed to characterize completely new host factor interactions, and strengthen observations from several previous studies.

**Author Summary:** The facultative intracellular pathogen *Salmonella enterica* serovar Typhimurium induces the reorganization of the endosomal system of mammalian host cells. This activity is dependent on translocated effector proteins of the pathogen. The host cells factors required for endosomal remodeling are only partially known. To identify such factors for formation and dynamics of endosomal compartments in *Salmonella*-infected cell, we performed a live cell imaging-based RNAi screen a to investigate the role of 496 mammalian proteins involved in cellular logistics. We identified that endosomal remodeling by intracellular *Salmonella* dependent on host factors in following functional classes: i) the late endo-/lysosomal SNARE (soluble *N*-ethylmaleimide-sensitive factor attachment protein receptor) complex, ii) the early secretory pathway, represented by regulators GTPases RAB1A and RAB1B, iii) the late secretory pathway and/or recycling endosomes represented by GTPases RAB3A, RAB8A, RAB8B, and the SNAREs VAMP2, VAMP3, and VAMP4, and iv) clathrin-coated structures. The identification of these new host factors provides further evidence for the complex manipulation of host cell transport functions by intracellular *Salmonella* and should enable detailed follow-up studies on the mechanisms involved.

## Introduction

The food-borne, facultative intracellular pathogen *Salmonella enterica* serovar Typhimurium (STM) is the etiological agent of gastroenteritis in humans or systemic infections in mice [1]. An early step in disease is the active invasion of epithelial cells. This process is dependent on the translocation of effector proteins by STM into the host cell through a type 3 secretion system (T3SS) encoded on *Salmonella* pathogenicity island 1 (SPI1) [2, 3].

After invasion STM, similar to many other intracellular pathogens, establish a replicative niche in host cells, termed *Salmonella*-containing vacuole (SCV). This process is dependent on the function of a distinct T3SS, encoded by SPI2 [4, 5] and translocating another set of effectors [6]. Though initially associating with markers of the early endosome (EE) such as EEA1 and the small GTPase RAB5 [7, 8], the SCV finally acquires several markers of the late endosome (LE). These include lysosome-associated membrane proteins (LAMPs) [9, 10], the vacuolar ATPase [11], and RAB7 [12, 13]. Concurrently, other canonical organelle markers such as the mannose-6-phosphate receptor are excluded [14].

A unique feature of STM among intravacuolar bacteria is the formation of a diverse array of long tubular structures, *Salmonella*-induced tubules (SIT) [15]. These include the LAMP-decorated *Salmonella*-induced filaments (SIF), the first SIT discovered [16, 17]. Moreover, SIF have been structurally characterized, revealing the presence of a double membrane tubular network [18, 19]. The host-derived membranes forming SCV, SIF, and other tubular compartments are collectively termed *Salmonella*-modified membranes (SMM).

The repertoire of bacterial effector proteins necessary for formation of SMM is quite well-characterized, with the SPI2-T3SS effector protein SifA being the most important factor [20, 21]. However, much less is known about corresponding host factors required for biogenesis of SMM. One crucial factor in SIF biogenesis is the SifA- and kinesin-interacting protein SKIP (a.k.a. PLEKHM2). In conjunction with the effectors SifA and PipB2 [22, 23] and the small GTPase ARL8B [24, 25], SKIP mediates kinesin-1 interaction and thus a link to the microtubule cytoskeleton and organelle motility [26].

Several attempts were made to analyze the interactions of STM with host factors in a systematic manner. These comprise RNA inference (RNAi) screens aiming at different parts of the STM infection process. Two genome-scale screens targeted the invasion [27, 28], while three screens focused on intracellular replication with two sub-genomic screens covering kinases and corresponding phosphatases, respectively [29, 30], and a genome-wide screen [31]. Additionally, two recent proteomic studies also shed light on interactions of intracellular STM with host cells. Vorwerk et al. [32] characterized the proteome of late SMM, while Santos et al. focused on early and maturing SCV [33].

All of these studies identified host factors yet unprecedented in STM pathobiology, and showed the general value of such systematic approaches. However, none of these approaches targeted specifically SIF, thus a host-SIF interactome is far from complete. Therefore, we established a targeted RNAi screen comprising 496 human genes mostly involved in cellular logistics to identify host factors involved in the formation of SIF. Using stably LAMP1-GFP-transfected HeLa cells, we performed automated microscopy on a spinning disk confocal microscope (SDCM) system with time-lapse live cell imaging (LCI) of STM infection, and scored for altered SIF formation as phenotypic readout. Investigating high-scoring hits of the RNAi screen, we validated several so far unknown host-SIF interactions by LCI: (i) involvement of the late endo-/lysosomal soluble *N*-ethylmaleimide-sensitive factor attachment protein receptor (SNARE) complex and its interaction partners, (ii) interactions of SIF with early secretory RAB1A/B, (iii) late secretory RAB3A, RAB8A/B, and VAMP2/3/4, and (iv) a connection to clathrin-coated structures.

## Results

### Setup and evaluation of RNAi screen

We aimed to identify host cell factors that are required for the endosomal remodeling induced by intracellular STM in a SPI2-T3SS-dependent manner. For this, an RNAi screen was performed with siRNAs predominantly targeting mammalian genes involved in cellular logistics and trafficking (75.2% categorized in intracellular transport according to Gene Ontology terms). This subset of 496 genes was termed ‘Trafficome’ and is listed in Table S1. Such a screen necessitates specific considerations and controls with the major ones described below, and further experimental issue are detailed in Suppl. Materials.

**Table 1.**
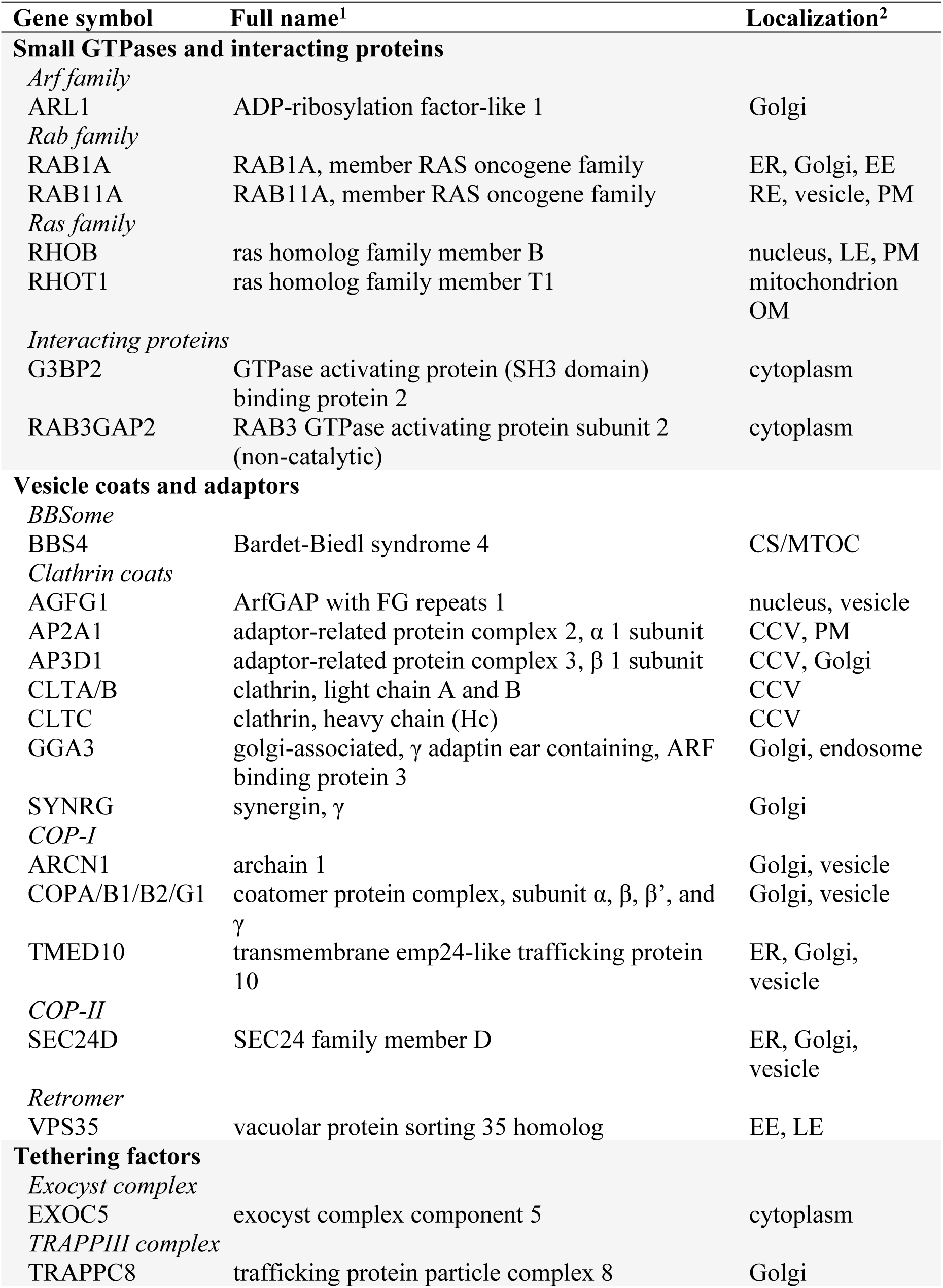
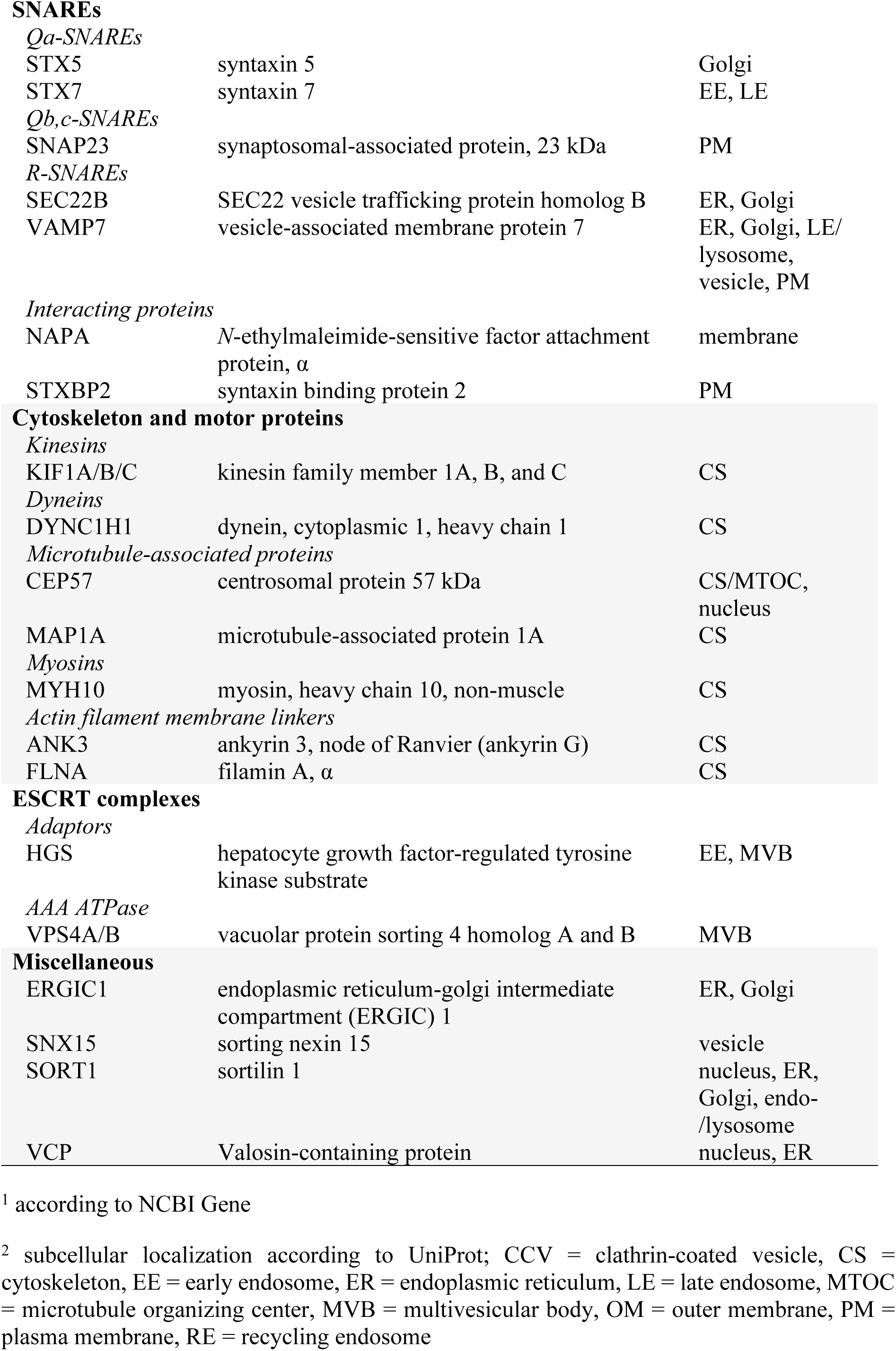
High-ranking trafficome hits (scoring cutoff of ≥8) involved in trafficking and cytoskeleton biology.

As a phenotypic readout for STM-induced endosomal remodeling, we scored the formation of SIF in infected cells. SIF show a highly dynamic behavior in their early phase after formation, with constant elongation and retraction [34, 35]. Thus, in contrast to previous RNAi screens done by analyzing fixed cells, we decided to perform this screen by LCI in order to obtain maximal phenotypic information. A previously established HeLa cell line stably transfected with LAMP1-GFP as the marker for SIF [18] was used as host cell.

As controls for STM-induced phenotypes, we used STM wild type (WT), capable in SIF induction, and an isogenic strain defective in SsaV, a central component of the SPI2-T3SS, and thus unable to induce SIF formation (Figure 1A). As a control for successful reverse transfection in general, we analyzed the lethal effect of an siRNA directed against polo-like kinase 1 (PLK1), a cell cycle control protein. The knockdown of this protein leads initially to a cell cycle arrest, and ultimately to cell death, as shown in Figure 1B. Besides, a phenotype-related control was established, i.e. knockdown of a host factor already known as essential for SIF formation. A host factor directly involved in SIF formation is SKIP [22]. This study already successfully used SKIP silencing, thus we used an siRNA with the same sequence as control. Real-time PCR indicated that the siRNA targeting SKIP yielded not a complete but sufficient and significant knockdown with a reduction to ca. 22% (Figure 1C). Transfection with AllStars siRNA did not affect SIF formation and dynamics over the course of infection (Figure 1D, Movie 1), while SKIP knockdown abolished SIF formation (Figure 1D, Movie 2) and reduced intracellular replication of STM. Though this does not completely exclude off-target effects, the phenotypic/visual control showed at least the intended purpose of this siRNA being fulfilled. The partial knockdown explains the rare appearance of SIF. Taken together, the establishment of the proper controls allowed us the execution of a larger scale RNAi screen.

**Figure 1.**
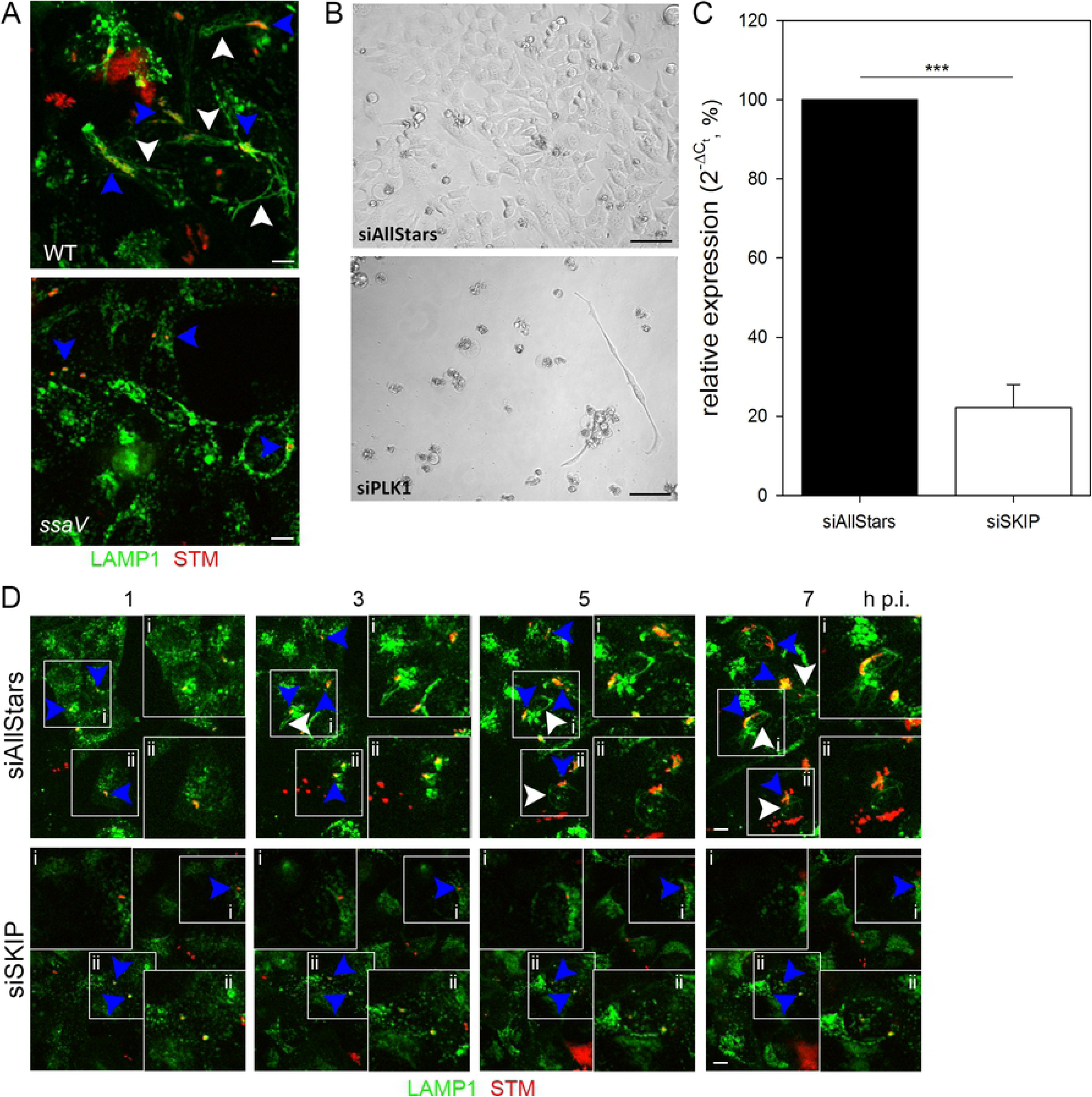
RNAi screen setup and validation. A) Intracellular phenotypes of STM under screening conditions. HeLa-LAMP1-GFP cells were infected with mCherry-labelled STM WT or *ssaV* strains and imaged live 8 h p.i. by SDCM. Presence of STM in LAMP1-positive SCV (blue arrowhead), induction of SIF formation by STM WT (white arrowhead), and lack for SIF formation by STM *ssaV* strain. Scale bar, 10 μm. B) Controls for siRNA-mediated knockdown. HeLa-LAMP1-GFP cells were reverse transfected with scrambled AllStars siRNA or PLK1 siRNA. Scale bar, 20 μm. C) Validation of SKIP siRNA knockdown. HeLa-LAMP1-GFP cells were reverse transfected with AllStars or SKIP siRNA. Then, RT-PCR targeting *SKIP* was performed. Depicted is the mean with standard deviation of three biological replicates (*n* = 3) each performed in triplicates. Statistical analysis was performed using Student’s *t*-test and indicated as: ***, *p* < 0.001. D) SKIP knockdown as control for inhibition of SIF formation. HeLa-LAMP1-GFP cells were first reverse transfected with AllStars or SKIP siRNA. Then, cells were infected with mCherry-labelled STM WT (MOI = 15) and imaged live 1-7 h p.i. by SDCM. Blue arrowheads indicate SIF-forming or non-SIF-forming single bacteria or microcolonies, white arrowheads indicate SIF. Scale bar, 10 µm.

### The RNAi trafficome screen

The complete workflow of the RNAi screen executed is summarized in Figure 2. First, siRNAs were automatically spotted onto 96-well plates. Additionally, each plate contained siAllStars, siPLK1, and siSKIP as negative and positive siRNA controls, and as phenotype-specific control, respectively. HeLa-LAMP1-GFP cells were seeded onto siRNAs for reverse transfection, incubated for 72 h, and subsequently infected with mCherry-labelled STM WT or *ssaV*. The formation of SIF was followed by LCI using an SDCM from 1-7 h post infection (p.i.) with hourly intervals.

**Figure 2.**
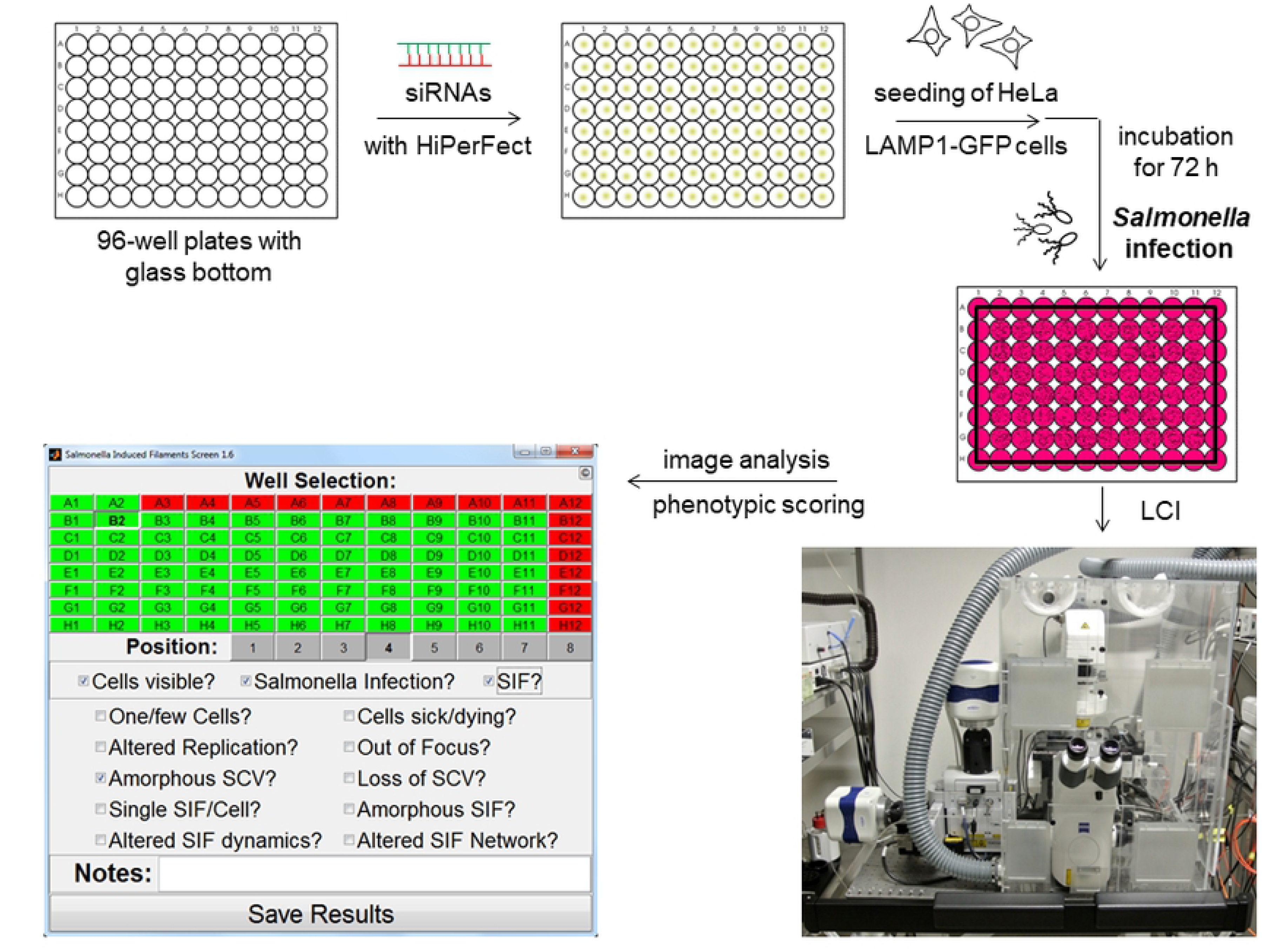
Basic workflow of the trafficome RNAi screen. 96-well plates with clear bottoms were automatically spotted with siRNAs. HeLa-LAMP1-GFP cells were seeded for reverse transfection. After 72 h of incubation, infection with STM was performed, followed by LCI using an SDCM system with hourly intervals of imaging. Phenotypic scoring was performed using the SifScreen utility. The MATLAB-based data input mask allows the entry of well- and position-specific information on general cell behavior and *Salmonella*/SMM phenotypes and the generation of a results report.

We set out to execute the analysis by visual inspection following the example of Stein et al. [20], who performed a mutant library screen to identify bacterial factors involved in SIF formation. As the dynamic nature of SIF and phenotypic heterogeneity in the cellular context excluded fully automated analysis, we decided to perform the analysis by visual inspections and used a MATLAB-based tool named SifScreen to support data input and collection (Figure 2). This tool queried the presence of SIF in the examined field of view as the main feature in a binary manner (for detailed information see Suppl. Material).

Since siRNA silencing usually does not yield 100% loss of function, we did not expect a complete lack of SIF in each of the eight images per well. Furthermore, since a considerable number of cells were present per image (roughly 10-30 cells, depending on applied siRNA and position on plate) a single SIF-forming cell would have prompted a SIF-positive scoring, even if generally a SIF-negative knockdown might have occurred. Thus, we decided to define an overall SIF-abolishing hit with a comparably high cutoff of 50%, i.e. if less than 50% of the images showed SIF. However, this did not take into consideration knockdowns possibly affecting cell viability in general or other circumstances compromising the analysis. Since these parameters were also queried by SifScreen, this allowed us to differentiate between ‘true hits’ and ‘possible hits,’ scoring the former and latter with values of 3 and 1, respectively. Additionally, to avoid a possible bias due to visual analysis, each screening plate was analyzed independently by two investigators. With the screen performed in biological triplicates, we subsequently compiled all scoring data for each host target, also pooling the results of the three individual siRNAs per target. This resulted in a list of final hits shown in Table S2, in which hits with a cumulative scoring of 1-4, 5-7, or ≥8 were classified as low-, mid- and high-ranking hits, respectively.

**Table 2.** Host proteins (gene symbols) identified as hits in the trafficome screen that are also present in at least one distinct SMM proteome.

| 8 h p.i. SMM<sup>1</sup> | 30 min p.i. SCV<sup>2</sup> | 3 h p.i. SCV<sup>2</sup> |
| --- | --- | --- |
| AP2A1 (AP-2) | - | AP2A1 |
| - | - | BET1 (SNARE) |
| - | CLTC (clathrin) | - |
| COPA/G1 (COPI)<sup>3</sup> | - | - |
| DYNC1H1 (dynein) | DYNC1H1 | - |
| - | ERGIC1 | ERGIC1 |
| - | ERP29 | - |
| - | - | EXOC5 |
| FLNA (filamin) | FLNA | FLNA |
| G3BP2 | - | G3BP2 |
| - | IQGAP1 | IQGAP1 |
| - | KIF5B (kinesin-1) | KIF5B |
| - | MAP1B | - |
| MYH9/10 (myosin II) | MYH9 | MYH9 |
| NAPA/α-SNAP | - | - |
| RAB2A<sup>3</sup> | RAB2A | - |
| - | RAB4A | - |
| RAB7A<sup>3</sup> | RAB7A | - |
| RAB11A<sup>3</sup> | - | - |
| RAB14<sup>3</sup> | - | - |
| - | SEC22B (SNARE) | - |
| - | SEC24C (COPII) | - |
| TMED10 (COPI) | - | - |
| VCP | - | - |

<sup>1</sup> [32]
<sup>2</sup> [33]
<sup>3</sup> colocalization with SMM shown by fluorescence microscopy

Approximately 81% (404 of 496) of the trafficome targets scored to varying degrees positive, underlining the general importance of trafficking processes for SIF formation. Table 1 shows selected high-ranking hits involved in trafficking and cytoskeleton biology. These hits clearly show the involvement of all protein classes necessary for the vesicle budding and fusion machinery, the core of cellular trafficking. These comprise: (i) small GTPases, especially Rab GTPases, as primary regulators [36–38]; (ii) vesicle coats and their adaptors as cargo and budding mediators [39–43]; (iii) cytoskeleton components as the basis for vesicle motility [44]; (iv) tethering factors as part of the fusion specification [45, 46]; (v) SNAREs as the primary fusion agents [45, 47, 48]. Besides, this list includes hits of diverse subcellular origin, encompassing the complete secretory and endo-/lysosomal system, i.e. endoplasmic reticulum (ER), Golgi apparatus, endo-/lysosomes. Supporting these allocations, the interaction network of the hits from Table 1 shows several distinct clusters (Figure 3). Two of them are connected to cytoskeleton biology (also interconnected if lower-ranking hits are included, data not shown). Another cluster is SNARE-centered, including RAB1A with RAB11A as a node between this cluster and one of the cytoskeleton-related clusters. Lastly, one cluster is associated with COPI and clathrin-coated vesicles (CCVs). Collectively, the overall results of the trafficome screen confirm the general importance of host trafficking factors in SIF biogenesis, indicating a crucial role for a plethora of as yet unprecedented factors in STM pathobiology.

**Figure 3.**
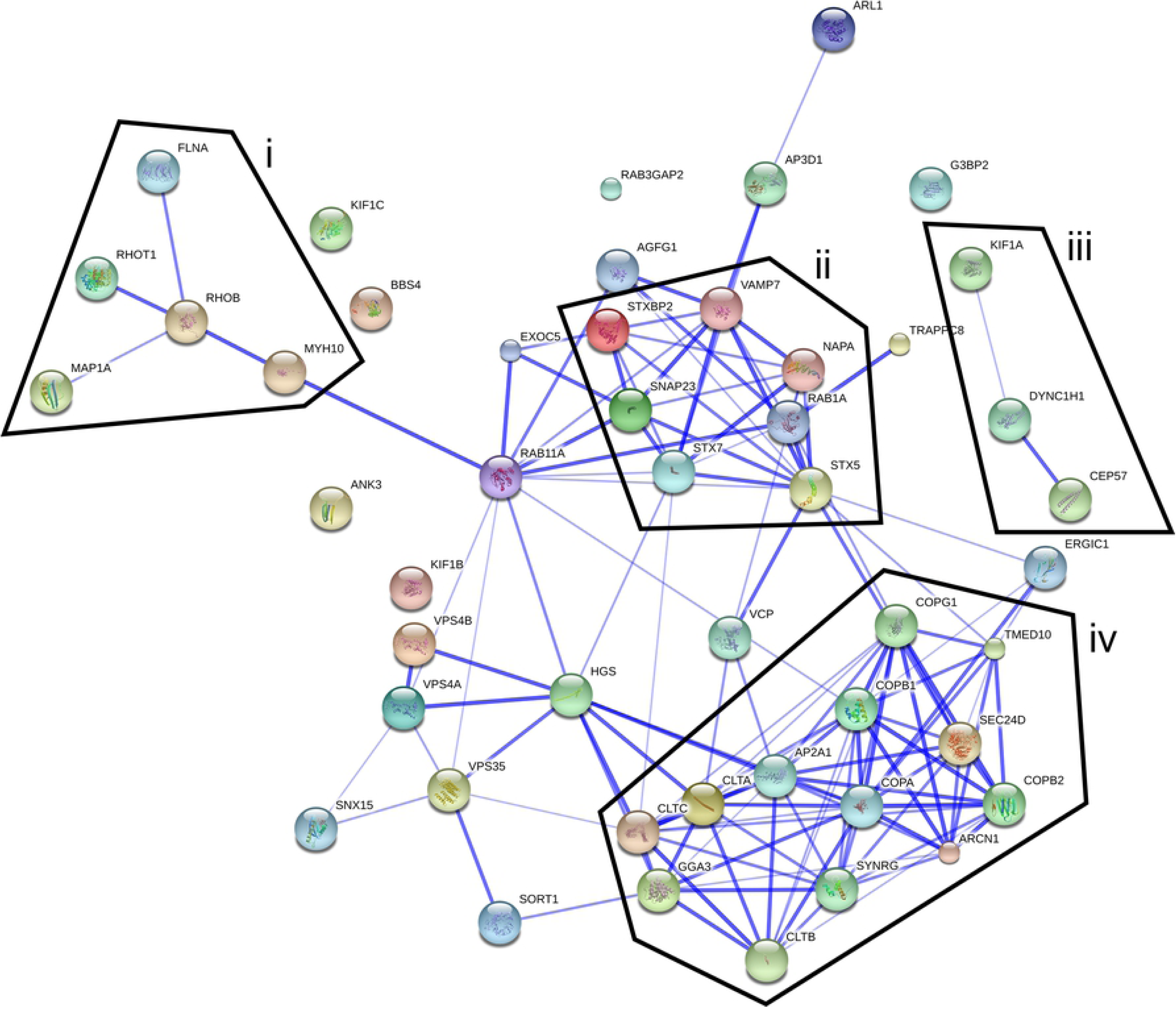
Interaction network of selected trafficome hits. The interaction of high-ranking hits (scoring cutoff of ≥8, see also Table 1) was visualized using the STRING database (confidence view). Borders delineate clusters related to the cytoskeleton (i, iii), SNAREs (ii), or COPI and clathrin-coated vesicles (iv).

### Validation of selected hits

To test the validity of our approach and the resulting hits, we focused on a subset of genes due to their presence in the noticeable interaction clusters depicted in Figure 3, or prior reports on involvement in STM pathobiology. HGS was chosen due to being the highest-ranking hit (Table S2) and the interaction of SPI1-T3SS effector SopB with endosomal sorting complex required for transport (ESCRT) complexes previously reported [49] with HGS being part of the ESCRT-0 complex. Furthermore, RAB1A and RAB11A were included as the highest-ranking Rab GTPases with RAB11A previously being shown to colocalize with SCV as well as SIF [32, 50]. Consistently, RAB7A served as another well-established SCV- and SIF-localizing control [12, 50–52]. STX5, STX7, VAMP7, and VAMP8 were chosen due to being the highest-ranking SNAREs (except VAMP8 lacking from the trafficome), the colocalization of STX7 with SIF [50], and the recent reported essential role of VAMP7 in SIF biogenesis [33]. The AAA ATPase VCP was included as another of the highest-ranking trafficking-related hits and another host factor already known to be important for proper SCV as well as SIF biogenesis via STM effector SptP [53]. Finally, the VPS11 core component of the class C core vacuole/endosome tethering (CORVET) / homotypic fusion and protein sorting (HOPS) group of multisubunit tethering complexes (MTCs) was chosen due to the HOPS complex functionally bridging RAB7 with late endo-/lysosomal SNAREs and the recent recognition of its essential role in STM replication and SCV and SIF biogenesis [54, 55].

The success of the siRNA knockdowns was confirmed by RT-PCR with a consistently significant decrease in mRNA in most cases down to 5-10% compared to control siAllStars (Figure S1, siSKIP served as the screen-inherent phenotype-specific control). Next, we exactly quantified the reduction of SIF formation due to the silencing of the selected targets (Figure 4). The *ssaV* mutant strain and the knockdown with siSKIP served as screen-inherent SIF-abolishing controls. All knockdowns resulted in decreased SIF formation with the reduction being statistically significant except for siHGS and siSTX5. The siRAB7A had the highest impact, siVCP the second-highest and the others ranged similar. Thus, the knockdowns of VAMP7 and VCP meet previous data (even though SIF abolishment regarding VCP depletion here is less pronounced) [33, 53]. Altogether, the effect on SIF formation demonstrates that our approach confirms known host factors, but also allows the identification of novel factors to be crucial for SIF biology.

**Figure 4.**
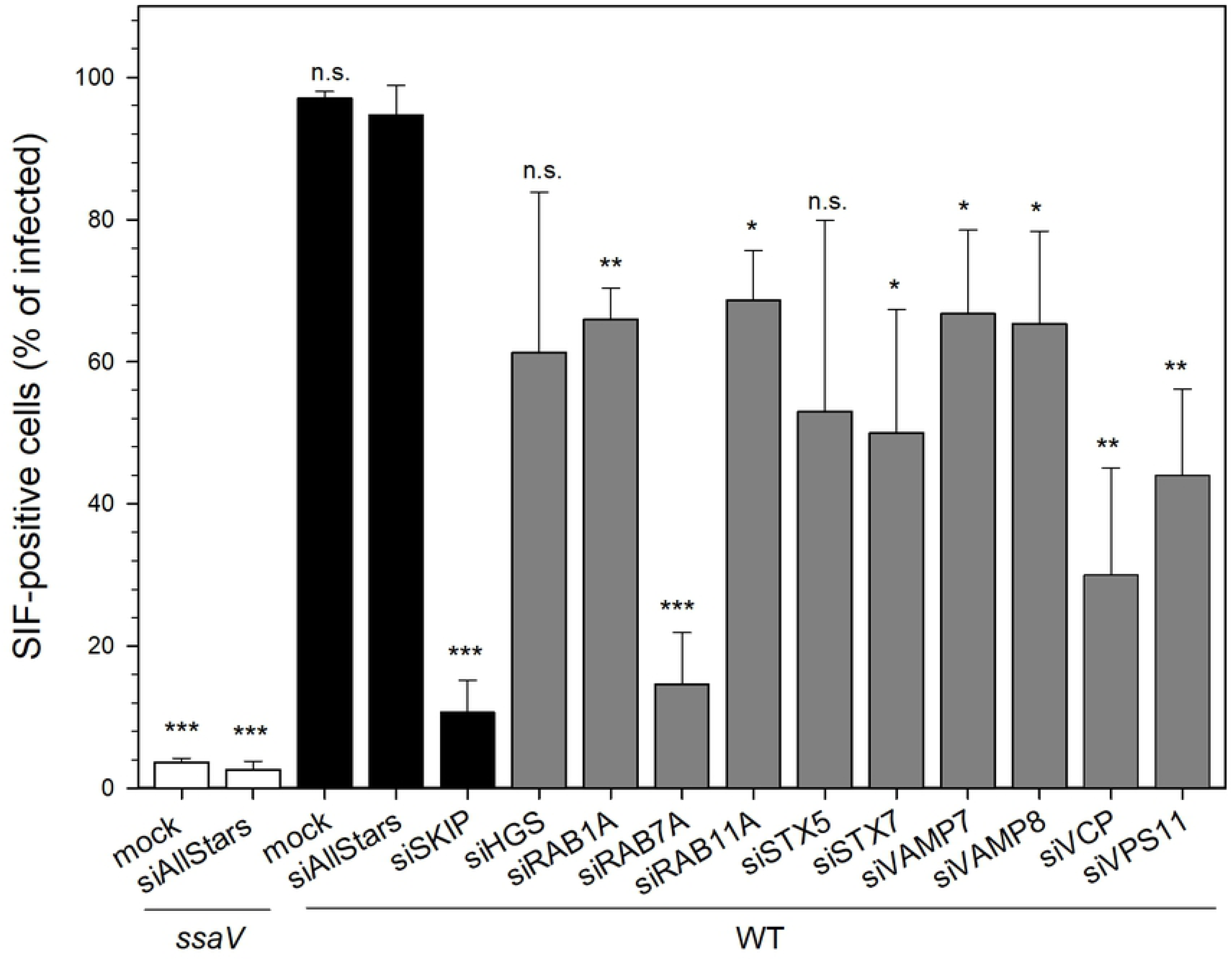
Influence of host factor silencing on SIF formation. HeLa-LAMP1-GFP cells were not transfected (mock), or reverse transfected with siAllStars or the indicated siRNA, infected with STM WT or SPI2-deficient *ssaV* expressing mCherry as indicated, and SIF counted. Depicted are means with standard deviation for three biological replicates (*n* = 3). Statistical analysis was performed against siAllstars + WT with Student’s *t*-test and indicated as: n.s., not significant; *, *p* < 0.05; **, *p* < 0.01; ***, *p* < 0.001.

### STM deploys membranes of early and late secretory, late endo-/lysosomal, and clathrin-coated origin in SIF biogenesis

The fact that host factors appear as hits in our screen, clearly indicates a physiologically relevant role in SIF biogenesis. However, whether this role is by direct interaction or an indirect one involving several intermittent steps, remains unclear. Thus, we decided to analyze the localization of selected hits with regard to SIF (Figure 5, Figure 6, Figure 7).

**Figure 5.**
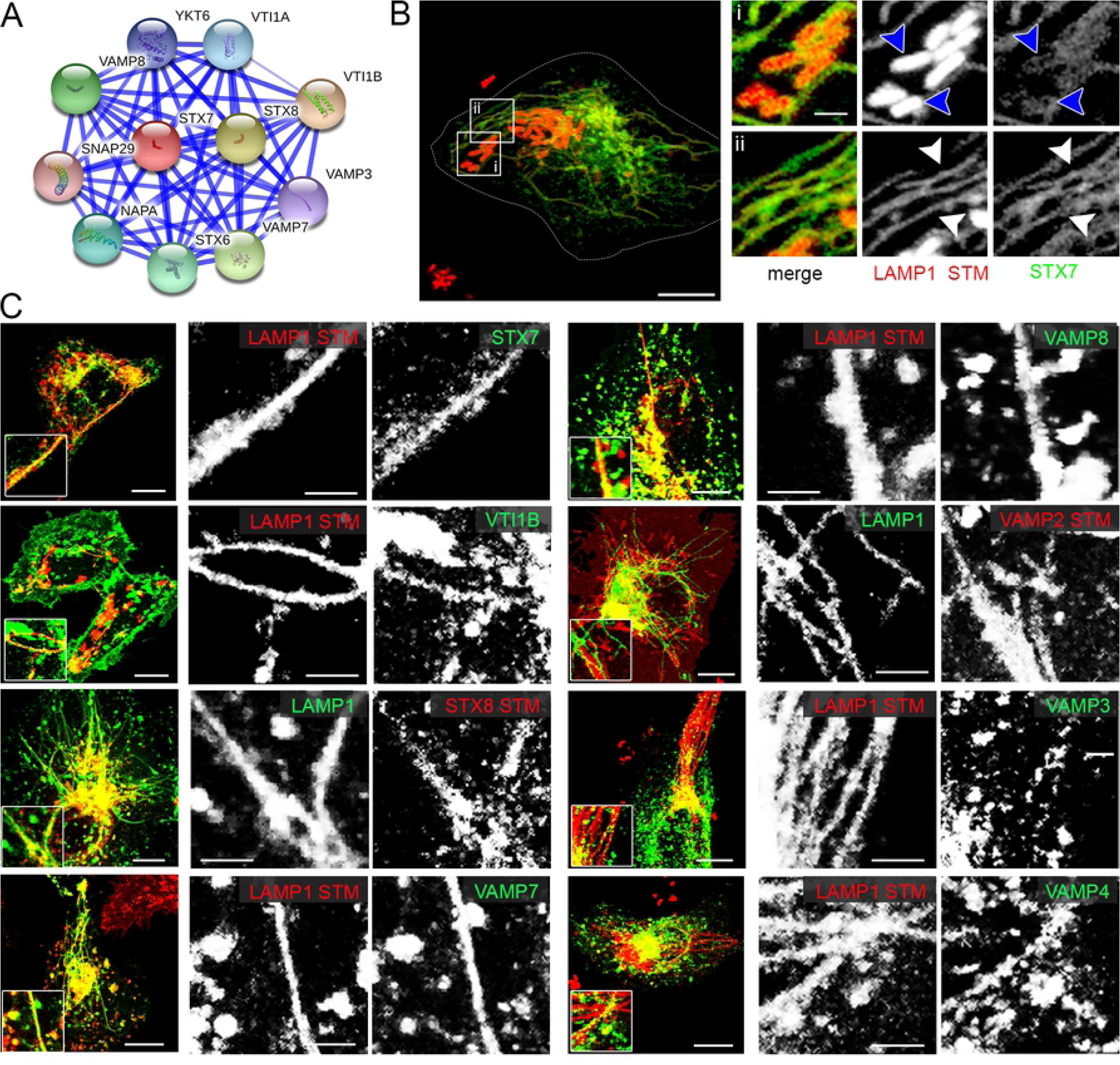
RAB proteins identified by the trafficome screen colocalize with SIF and SCV. A) Direct interaction network of RAB1B as visualized by STRING. B) and C) HeLa cells either stably transfected with LAMP1-GFP (green), or transiently transfected with LAMP1-mCherry (red) were co-transfected with plasmids encoding various RAB GTPases (RAB7A, RAB1A, RAB1B, RAB3A, RAB8A, RAB8B, RAB9A, RAB11A) fused to GFP (green) or mRuby2 (red) and then infected with STM WT expressing mCherry or GFP. Living cells were imaged from 6-9 h p.i. by CLSM and images are shown as maximum intensity projections (MIP). Insets magnify structures of interest and white arrowheads indicate colocalization with SIF. Scale bars, 10 µm (overviews), 1 µm (details).

**Figure 6.**
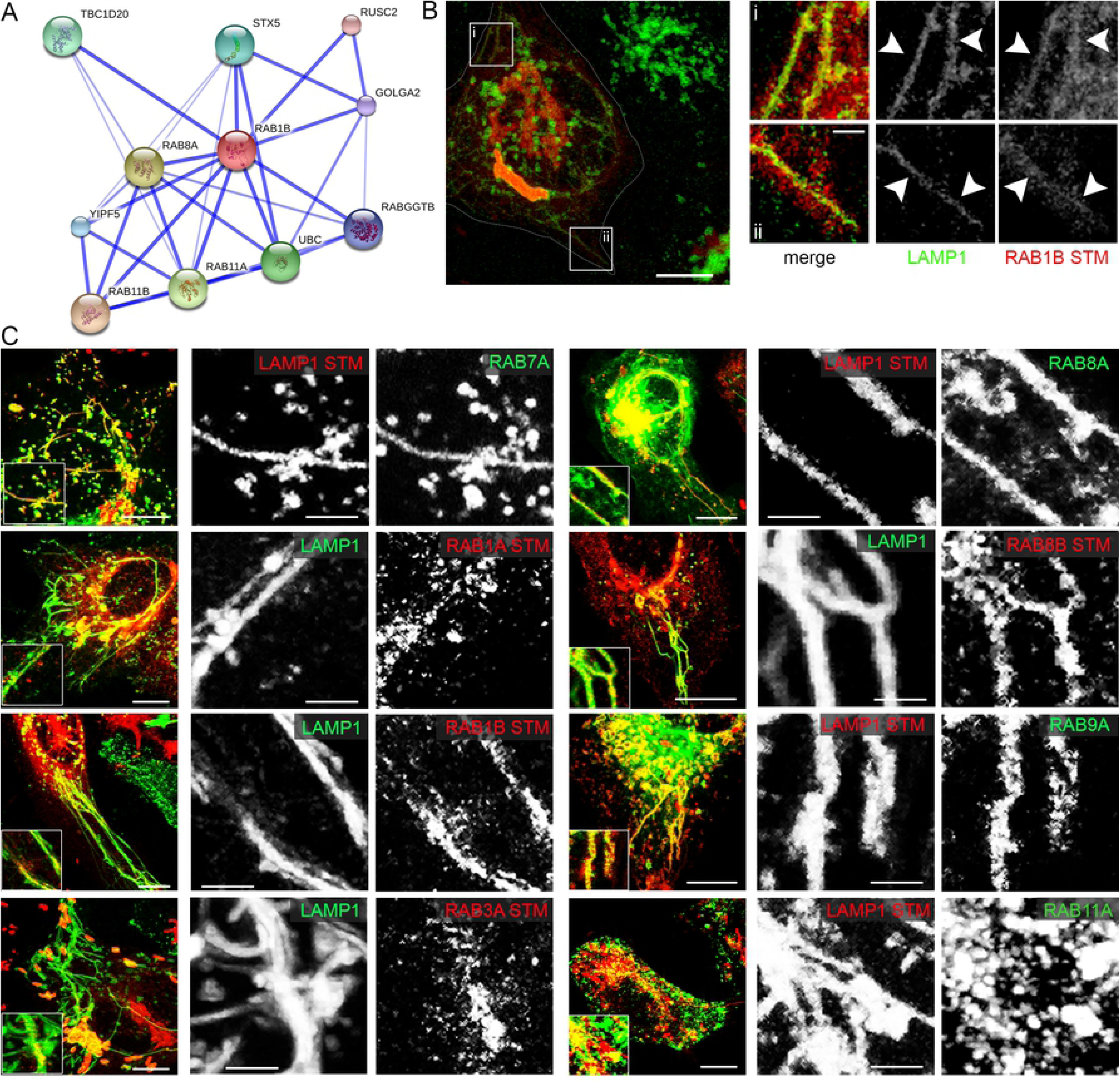
SNARE proteins identified by the trafficome screen colocalize with SIF and SCV. A) Direct interaction network of STX7 as visualized by STRING. B) and C) HeLa cells either stably transfected with LAMP1-GFP (green), or transiently transfected with LAMP1-mCherry (red) were co-transfected with plasmids encoding various SNAREs (STX7, VTIB, STX8, VAMP7, VAMP8, VAMP2, VAMP3, VAMP4) fused to GFP (green) or mRuby2 (red). Infection and imaging was performed as for Figure 5. Insets magnify structures of interest and white and blue arrowheads indicate colocalization with SIF and SCV, respectively. Scale bars, 10 µm (overviews), 1 µm (details).

**Figure 7.**
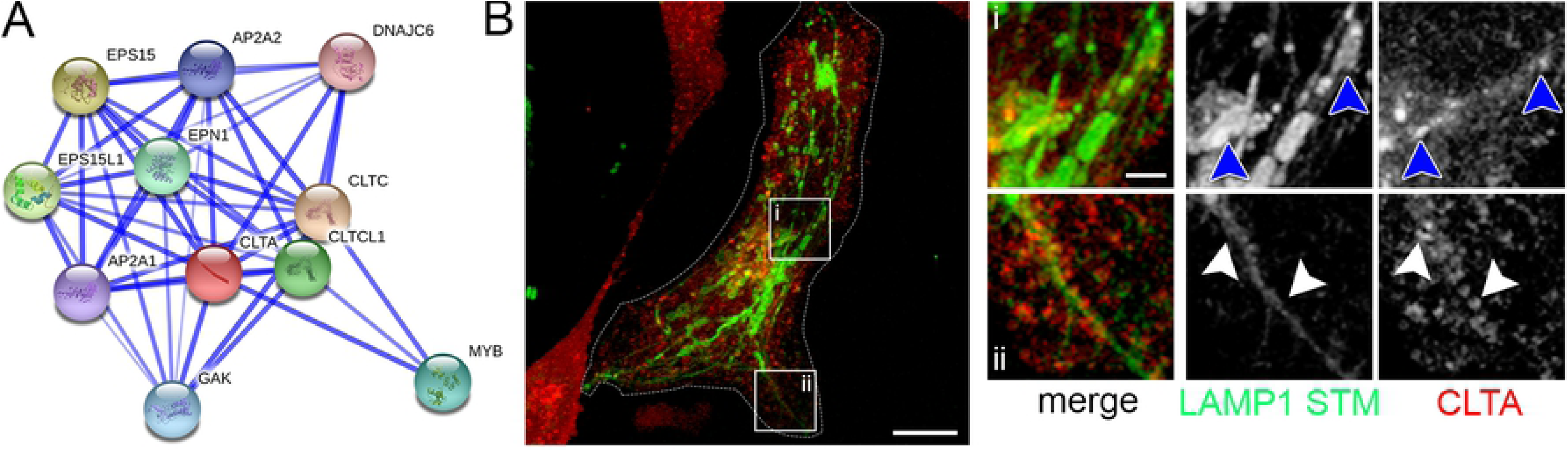
CLTA identified by the trafficome screen colocalizes with SIF and SCV. A) Direct interaction network of CLTA as visualized by STRING. B) HeLa cells stably transfected with LAMP1-GFP (green) were co-transfected with a plasmid encoding CLTA fused to mRuby2 (red). Infection and imaging was performed as for Figure 5. Insets magnify structures of interest and white and blue arrowheads indicate colocalization with SIF and SCV, respectively. Scale bars, 10 µm (overviews), 1 µm (details).

For analyses of RAB GTPases (Figure 5), we again used RAB7A as well as RAB9A as positive controls, both showing a clear colocalization with SIF. Of the several Rab GTPases included in the trafficome RAB1A showed the highest score (Table S2). RAB1 GTPases are responsible for anterograde ER-Golgi trafficking [56–59]. Importantly, RAB1A can be functionally substituted by RAB1B [60, 61] and an STM replication-targeted RNAi screen identified specifically RAB1B as a hit [31]. Hence, we analyzed the infection-related localization of both isoforms and detected a partial and a strong colocalization of RAB1A and RAB1B, respectively, with SIF (Figure 5).

Another high-ranking hit with relation to RAB proteins was RAB3GAP2 (Table S2), the non-catalytic subunit of the RAB3 inactivating GTPase-activating protein (GAP) complex [62]. RAB3 possesses four isoforms in mammals [63] and is involved in regulated exocytosis [64]. As neither the catalytic GAP subunit, RAB3GAP1, nor one of the four isoforms were present in the trafficome, we decided to analyze the localization of RAB3A and found a partial colocalization with SIF (Figure 5).

Besides, a mid-ranking hit was RAB8A (Table S2), a Golgi- and endosome-localized RAB likewise involved in exocytic processes [65]. Interestingly, its isoform RAB8B was observed to be excluded from maturing SCVs (≤ 3 h p.i.) [50]. Therefore, we analyzed the localization of both, RAB8A and RAB8B, and strikingly found a strong colocalization of not only RAB8A but also RAB8B with SIF (Figure 5).

As several RAB proteins participating in the late secretory system/exocytosis seem to play a role in SIF biogenesis, we additionally analyzed three SNAREs with exocytic roles not present in the trafficome: VAMP2, VAMP3, and VAMP4 [66–68], with VAMP2 also shown to be present on early SCV [69]. Apart from that, the presence of the two high-ranking SNARE hits STX7 and VAMP7 (Table S2) on SIF was previously shown [33, 50]. However, SNAREs, are part of complexes of usually four proteins participating in membrane fusion and consisting of a single protein v-SNARE (on the vesicle or incoming membrane) and a ternary t-SNARE subcomplex (on the target or accepting membrane). VAMP7 is the v-SNARE in the SNARE complex for heterotypic LE/lysosome fusions with the t-SNAREs STX7, STX8, and VTI1B [70, 71] being replaced by VAMP8 in homotypic LE fusions [72, 73]. In fact, the presence of VTI1B and STX8 on early SCV [69, 74] and their role in STM replication [54], as well as the involvement of VAMP8 in STM invasion were already shown [75]. However, their interaction with SIF remains unclear, except VAMP8 silencing causing SIF reduction identified here (Figure 4). Thus, we analyzed the localization of VTI1B, STX8, and VAMP8 using STX7 and VAMP7 as controls. As shown in Figure 6, we detected prominent association of STX7 and VAMP7 with SIF, as well as for VAMP2 and VAMP8. Colocalization of VTIB, STX8, VAMP3, and VAMP4 with SIF was also observed. However, these SNARE subunits showed a more heterogeneous distribution, and only a fraction of SIF was positive for these candidates. Notably, both recent proteomic studies [32, 33] and this screen identified AP2A1 as being present on SMM/SCV or scoring high-ranking (Table S2), respectively. AP2A1 represents one of the two core subunit isoforms of the canonical AP-2 adaptor complex usually acting in clathrin-mediated endocytosis (CME) at the plasma membrane [76, 77]. Accordingly, the main coat determinants implicated in the formation of CCVs were among the high-scoring hits, including both clathrin light chains, CLTA and CLTB, as well as the conventional heavy chain, CLTC (Table S2). Thus, we analyzed the localization of CLTA and observed a partial colocalization with SIF (Figure 7).

In conclusion, the colocalization of various host factors involved in cellular transport with SIF validated the results of the RNAi screen. These proteins are components of SIF tubules and, to variable extent, required for the formation of SIF. The data also support the highly diverse origin of host cell membranes involved in SIF formation.

## Discussion

By applying a targeted RNAi screen, we identified several new host factors required for the formation of SIF, and partially characterized interactions of host proteins with SMM. Out data strengthen the involvement of the late endo-/lysosomal SNARE complex, and reveal new interactions of SIF with RAB1, RAB3, and RAB8 GTPases, exocytic SNAREs, as well as clathrin-coated structures. The implications of these findings as discussed below are depicted in Figure 8. Several host trafficome components identified here were previously shown as involved in infection biology in STM in general, and specifically in SCV and/or SIF biogenesis including: dynein – DYNC1H1 [78–80], filamin – FLNA [81], myosin II – MYH10 [82], VPS4A/B [49]. This also holds true for several mid-ranking hits: kinesin-1 – KIF5A/B [22–25, 83], PIKFYVE [84], RAB9A [85, 86], RAB14 [85], SCAMP3 [87].

**Figure 8.**
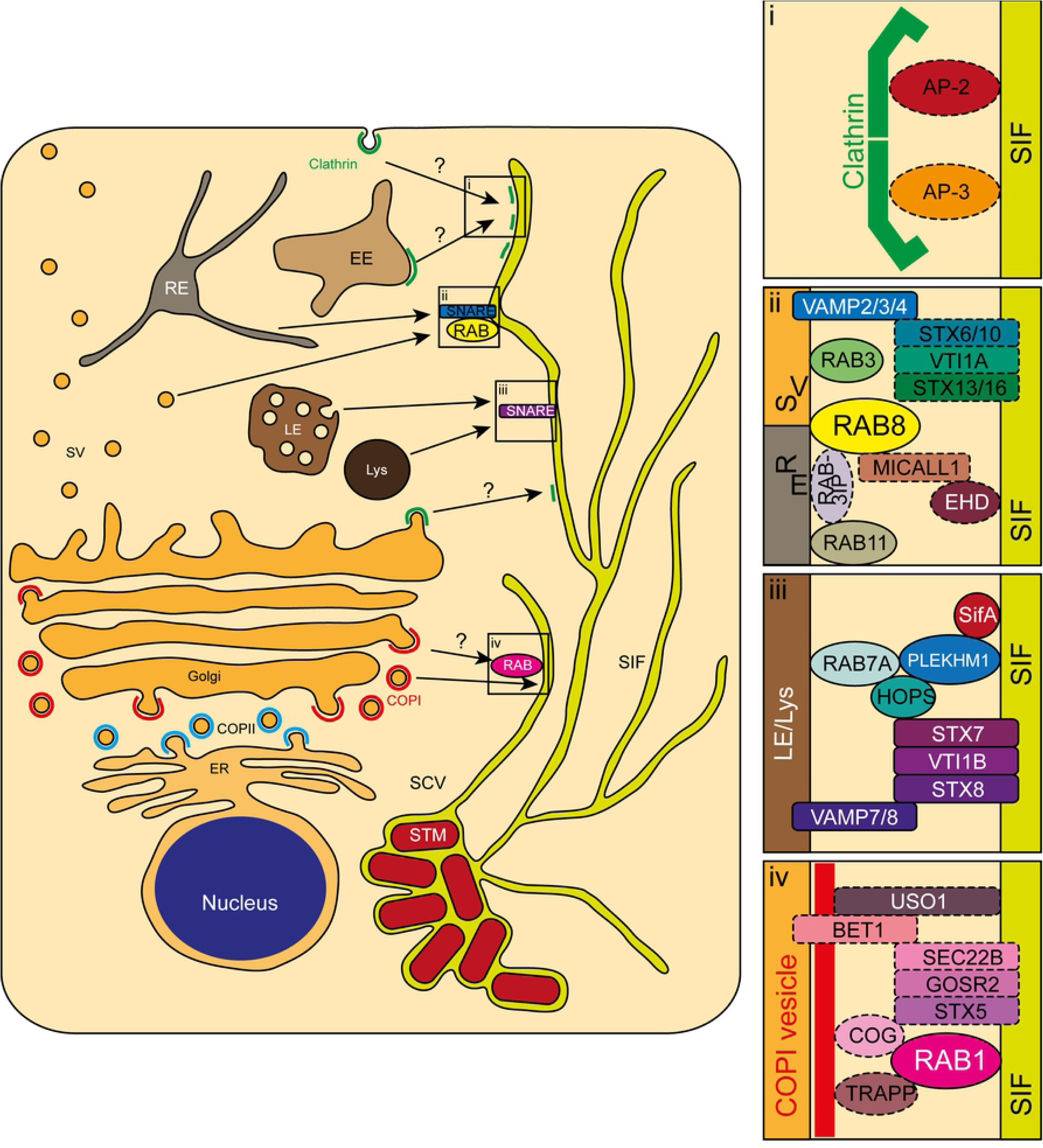
Newly identified interactions of intracellular STM with host factors. Depicted are central eukaryotic endomembrane organelles possibly playing a role in the newly identified interplays of host factors with SIF. Magnifications show the interactions of clathrin (i), late secretory and/or recycling-related RAB3A, RAB8A/B, and VAMP2/3/4 (ii), late endo-/lysosomal VTI1B, STX8, and VAMP8 (iii), and early secretory RAB1A/B (iv) with other host factors added as discussed in the text. Solid lines represent interactions identified here or otherwise known, dashed lines represent putative interactions. COP, coat protein complex; EE, early endosome; ER, endoplasmic reticulum; LE, late endosome; Lys, lysosome; RE, recycling endosome; SCV, *Salmonella*-containing vacuole; SIF, *Salmonella*-induced filaments; STM, *S.* Typhimurium; SV, secretory vesicle.

Complementary data were recently provided by two proteomic studies. Our group analyzed the SMM proteome in the late phase of infection (8 h p.i.) [32] that contained several host proteins that are mid- or high-ranking hits in this screen (summarized in Table 2, first column). The colocalization of several of these proteins with SMM was shown by immunostaining or LCI. Santos et al. [33] determined the proteomes of early and maturing SCV (30 min p.i. and 3 h p.i., respectively) again identifying proteins appearing as hits in this screen (see Table 2, second and third column). Taken together, these data strongly validate the approach deployed here.

The approach reported were has an major advantage compared to studies based on organelle proteomics [32, 33]. Proteomics shown presence or absence of host factors on the organelle of interest, but a particular role in the biogenesis of this organelle cannot be implied directly. In our RNAi approach potentially each, or at least each high-ranking hit, points to a role in STM-induced endosomal remodeling. However, a functional role revealed by RNAi does not necessarily require colocalization of the host factor with the compartment, because a function may be mediated indirectly, involving several interacting partners. We analyzed the localization of selected factors (Figure 5, Figure 6, Figure 7) and found several differences in the host factor sets identified by proteomics or by our approach. Nevertheless, there is a considerable overlap of host factors identified by both approaches as represented in Table 2.

The interaction of microbial pathogens with regulators of the early secretory system, such as *Legionella* and *Coxiella* with RAB1 [88–90], and *Brucella* with RAB2 [91, 92], is well established. However, an interaction of STM with the early secretory system, e.g. RAB2A, was only recently described by proteomic studies [32, 33]. We now expand this interaction by showing the physiological relevance of RAB1A in SIF formation (Figure 4), as well as the presence of both RAB1A and RAB1B on SIF (Figure 5). This is in striking contrast to the previous observation that RAB1A is detrimental to STM replication due to its role in antibacterial autophagy, which is counteracted by the SPI2-T3SS effectors SseFG [93, 94]. Future work has to clarify this apparent conundrum, at least regarding RAB1A. The direct association of RAB1A/B with SIF possibly connects several distinct trafficking events. First, RAB1B was shown to be involved in formation of the COPI vesicle coat, which participates in *intra*-Golgi and retrograde Golgi-to-ER transport. The formation of COPI is dependent on RAB1B due to its effector GBF1, which is an activating guanine exchange factor (GEF) of ARF1, the primary COPI-regulating small GTPase [95–97]. Second, the COPI components COPA and COPG1 were shown to partly colocalize with SIF [32]. In accordance, our screen identified the majority of COPI components as mid- or high-ranking hits (ARCN1, COPA/B1/B2/G1, Table 1 and Figure 3, though ARF1 did not score at all, and GBF1 only low). Thus, RAB1A and/or RAB1B might represent a physical link between COPI vesicles and SCV and/or SIF for the redirection of early secretory material as depicted in Figure 8iv. The physical interaction of SIF with COPI vesicles might be, similar to the late endo-/lysosomal fusion machinery, additionally accompanied by tethering factors and SNAREs. The conserved oligomeric Golgi (COG) tethering complex was shown to be a RAB1 effector and directly bind COPI components [98, 99]. Interestingly, all components of the COG present in the trafficome scored mid- to high-ranking (COG1/2/3/5/7, Table S2). Additionally, COG binds STX5, a SNARE that is part of several ER-Golgi and *intra*-Golgi transport-related SNARE complexes [100]. These complexes either consist of STX5, GOSR2/GS27/membrin, BET1, SEC22B [101], or STX5, GOSR1/GS28, BET1, YKT6 [102], or STX5, GOSR1, BET1L/GS15, YKT6 [103]. All of these SNAREs are present in the trafficome, but strikingly only the components of the first SNARE complex scored all mid- to high-ranking (Table 1 and Table S2). STM effectors partaking in this interaction might by SseF and SseG, as they were recently shown in a BioID screen [104] to interact with STX5 and SEC22B, besides PipB2 also interacting with SEC22B. However, the potential involvements indicated by these collective data remain to be elucidated.

Transport protein particle (TRAPP) complexes I, II, and III were identified as RAB1 GEFs [105–108], as well as COPII [TRAPPI; 109, 110, 111], and COPI tethers [TRAPPII; 107, 112]. TRAPPI is the core shared by all TRAPP complexes, with II and III processing unique additional subunits. TRAPPC8, the unique component of TRAPPIII, scored high-ranking. Other components were not present in this trafficome, except the TRAPPI core subunit TRAPPC2, which unexpectedly did not score at all (Table S2). So far TRAPPIII is only characterized to participate in autophagy [108, 113, 114]. Finally, the golgin USO1/p115 scored mid-ranking (Table S2). It is also a RAB1B effector and COPI and COPII tether [97, 115, 116], partly in conjunction with COG [117], besides being likewise able to bind STX5 [118]. For both TRAPP complexes and USO1, a specific role in SIF biogenesis remains to be elucidated. It has already been described that STM, depending on SseF/SseG, recruits to the SCV exocytic vesicles from the Golgi apparatus destined to the plasma membrane [119]. Which host factors are involved in this process was unclear, and our work now sheds light on this phenotype by showing the presence of exocytic RABs, RAB3A, RAB8A, and RAB8B (Figure 5) on SIF, as well as exocytic SNAREs, i.e. VAMP2, VAMP3, and VAMP4 (Figure 6).

Besides their involvement in exocytosis, VAMP4 and VAMP3 are also known to prominently participate in endosome-to-Golgi transport in conjunction with STX16, VTI1A, and STX6 or STX10 for EEs or LEs, respectively [120, 121]. Although STX6 and STX10 were not included in the trafficome and STX16 ranked low, VTI1A was a mid-ranking hit (Table S2). The STM-mediated redirection of LAMP1-containing vesicles from the Golgi apparatus to the early SCV was shown to involve recruitment of STX6 and VAMP2 via SPI1-T3SS effector SipC [69]. Alternatively, this might happen via SPI2-T3SS effector PipB2 that was identified in the recent BioID screen as interactor of VAMP2 [104]. Furthermore, in homotypic EE fusion STX16 is replaced by STX13 [122], and STX13 was previously shown to be present on early SCV [74, 123]. While the exact role of RAB3A and the identity of SNAREs involved remain to be determined, this might indicate that the interception of secretory vesicles depends on a SNARE complex comprising a distinct combination of the abovementioned SNAREs as represented in Figure 8ii.

In addition to its exocytic role, RAB8A is involved in recycling processes as indicated by the localization on tubular recycling endosomes (RE) [124]. This localization depends on several factors such as RAB8 GEF RAB3IP/RABIN8 (which is also part of the trafficome, though it ranked low, Table S2), concurrently being an effector of RE master regulator RAB11 [125]. Another factor is the RE-localized MICALL1, which interacts with the dynamin-like ATPases EHD1 and EHD3 [124, 126, 127]. Interestingly, MICALL1 was identified in an RNAi screen with focus on STM intracellular replication [31], EHD1 and EHD2 were present in the proteome of maturing SCV [33], and EHD4 was present in late SMM [32]. Moreover, the association of the maturing SCV and late SMM with RAB11A/B was shown previously [32, 50], with RAB11A following RAB1A as the second highest-ranking RAB in our screen (Table S2). RAB8B was not present in our screen, but showed a clear colocalization with SIF. In a previous screen RAB8B was shown to be excluded from maturing SCV [50], however only early events up to 3 h p.i. were analyzed. Temporal differences in RAB8B recruitment could explain these observations. SPI2-T3SS effector SopD2 most likely plays a role in RAB8 recruitment, as it was previously shown to interact with RAB8 [104, 128]. Collectively, these data strongly argue for a continued association of STM not only with exocytic compartments, but also with recycling compartments at later time points (summarized in Figure 8ii). Concurrently, a RAB11-to-RAB8 switch might occur on SMM, similar to the RAB5-to-RAB7 and RAB14-to-RAB9 switches, probably involving RAB3IP/RABIN8. However, unlike the enduring RAB5-to-RAB7 switch and similar to the RAB14-to-RAB9 switch, this seems to happen repeatedly in a transient manner due to the continued presence of RAB11 on SIF as with RAB14 [32].

The presence on SIF, and/or importance for SIF formation of RAB7, the HOPS complex, STX7, and VAMP7, as well as the direct fusion of late endo-/lysosomal-like VAMP7-positive vesicles with the SCV, was shown before [33, 50, 54]. This indicates the involvement of the complete canonical mammalian late endo-/lysosomal vesicle fusion machinery in SIF biogenesis. Whether this interaction cascade also employs the canonical STX7, VTI1B, and STX8 was not fully clarified. Here, we expand this cascade by showing the physiological relevance of STX7 for SIF formation (Figure 4), and the presence of VTI1B and STX8 on SIF (Figure 6, Figure 5) as depicted in Figure 8iii. This cascade is possibly expanded by the host protein PLEKHM1, as the recruitment of RAB7 and the HOPS complex by SifA via the host protein PLEKHM1 and its involvement in SCV biogenesis was recently revealed [55], most likely also being involved in SIF biogenesis. Taken together, SifA seems to recruit the complete late endo-/lysosomal fusion machinery. Thus, SifA performs a dual role besides the binding of SKIP and the SIF mobility connected with it. This is also corroborated by the identification of interactions of SifA with STX7 and VAMP7 by the recent BioID screen [104]. Alternatively or in addition, SopD2 might be likewise involved as it was also shown to interact with STX7 and VAMP7 besides VTI1B in the same study.

Data on involvement of clathrin-coated structures or adaptor protein complexes in infection processes of intracellular bacterial pathogens are scarce. The usurpation of CME during the internalization/invasion process is only recently being recognized to be employed by several bacteria, best studied in *Listeria monocytogenes* infection [129, 130]. However, only one study on *Brucella abortus*, a pathogen also residing in a vacuole during intracellular lifestyle, demonstrated the association of clathrin with the *Brucella*-containing vacuole [131]. We now show such an association with CLTA also for STM (Figure 7). It is peculiar that proteomics, as well as our screen, indicate an involvement of the AP-2 complex, but one of the other adaptor complexes. This is noteworthy due to the CME-related AP-2 being primarily plasma membrane-localized, in contrast to the Golgi traffic-related AP-1 and AP-4, or the endo-/lysosomal traffic-related AP-3 and AP-5 [43]. Especially AP-3 deserves detailed analyses since its two core subunit isoforms scored in mid- and high-ranking range (AP3B1 and AP3D1, Table S2, see Figure 8i). An interaction of AP-3 with VAMP7 in mammalian cells [132–134], as well as an interaction with the complete late endo-/lysosomal SNARE complex in *Dictyostelium discoideum* was previously shown [135]. Several SPI2-T3SS effectors, i.e. PipB2, SopD2, and SseG, might participate in such a recruitment because a recent BioID screen revealed interaction with various AP-2 and AP-3 core subunits [104]. However, examination of other AP complexes also seems worthwhile, since the latter study indicates interactions with several of them and the trafficome screen did not comprehensively cover AP complex.

In summary, we successfully employed a sub-genomic RNAi screen to systematically identify new host factors, corresponding protein complexes, and pathways involved in SIF formation. By providing physiologically relevant data regarding SIF formation, this work further corroborates the promiscuous origin of SMM indicated by previous proteomics studies [32, 33]. Similar future screens can also reveal the biogenesis of several other SIT [15], and extend to the host cell types important for *Salmonella* pathogenesis.

## Acknowledgements

We thank Monika Nietschke and Ursula Krehe for construction of plasmids and technical assistance. Special thanks go to Markus C. Kerr (Brisbane), André P. Mäurer, and Thomas F. Meyer (Berlin) for advice and logistics in setting up the screen. We thank Martin Aepfelbacher (Hamburg), Thierry Galli (Paris), Wanjin Hong (Singapore), and Yulong Li (Beijing) for providing transfection vectors, and acknowledge DNASU und Addgene for provision of materials.

## Materials and Methods

### Bacterial strains and growth conditions

For infection STM NCTC 12023 WT and isogenic SPI2-T3SS-defective strain P2D6 harboring plasmid pFPV-mCherry/2 or isogenic GFP-expressing MvP1897 were used (for details see Table S6). Strains were routinely grown in Luria-Bertani (LB) broth (Difco, BD, Heidelberg, Germany) containing 50 µg/mL carbenicillin for plasmid selection at 37 °C with aeration.

### Cell lines and cell culture

Experiments were performed using the parental HeLa cell line (ATCC No. CCL-2) or the lentivirus-transfected HeLa cell line stably expressing LAMP1-GFP [18]. Cells were routinely cultured in Dulbecco’s modified Eagle’s medium (DMEM) containing 4.5 g/L glucose, 4 mM stable glutamine, and sodium pyruvate (Biochrom, Berlin, Germany) supplemented with 10% inactivated fetal calf serum (iFCS; Gibco, Darmstadt, Germany) in an atmosphere of 5% CO_2_ and 90% humidity at 37 °C.

### siRNA library and individual siRNAs

The siRNA library used comprised siRNAs targeting 496 host proteins mostly involved in intracellular trafficking (but also other processes such as metabolism) with a threefold coverage. The siRNAs were part of a human whole-genome library obtained from Qiagen (Hilden, Germany) deposited at the Max Planck Institute for Infection Biology (Berlin, Germany). A volume of 4 µL of each siRNA (0.2 µM, end concentration of 5.2 nM) was spotted automatically onto 96-well Clear Bottom Black Cell Culture Microplates (Corning, Corning, NY, USA) and frozen at -20 °C before transfer. Additionally, each plate contained the same amount of the following siRNAs from Qiagen as knockdown controls: AllStars as negative and Hs_PLK1_7 (directed against the cell cycle protein polo-like kinase 1) as positive controls. A custom siRNA from Qiagen directed against SKIP served as a phenotype-specific control [22] and was spotted on location. Information including target sequences for these siRNAs, as well as those ordered for validation experiments, are listed in Table S7.

### Reverse transfection with siRNA

If not using 96-well screening plates as detailed above, the amount for an end concentration of 5 nM siRNA was spotted onto standard cell culture 6-well plates (for mRNA extraction, TPP, Trasadingen, Switzerland) or 8-well polymer bottom chamber slides (for quantification of SIF formation, µ-Slides, ibidi, Martinsried, Germany).

Next, a mixture of the transfection reagent HiPerFect (Qiagen, Hilden, Germany) and serum-free cell culture medium was applied and this was incubated for 5-10 min at room temperature (RT). Subsequently, 5,000, 125,000, or 20,000 cells per well of 96-well plates, 6-well plates, or 8-well chamber slides, respectively, were added in serum-containing medium and incubated for 72 h at 37 °C in a humidified atmosphere containing 5% CO_2_.

### Gene expression quantification

After reverse transfection with different siRNAs total RNA of cells was extracted using the RNeasy Mini Kit following the manufacturer’s instructions (Qiagen, Hilden, Germany). Homogenization during extraction was performed using Qiagen QIAshredder columns. Then, 1 µg of RNA digested with DNaseI (NEB, Frankfurt a. M., Germany) was used for reverse transcription of mRNA with the RevertAid First Strand cDNA Synthesis Kit (Thermo Scientific, Dreieich, Germany) following the manufacturer’s instructions using the Oligo(dT)_18_ primer. For RT-PCR 1 µL of cDNA was used with the Thermo Scientific Maxima SYBR Green/Fluorescein qPCR Master Mix (2x). As reference gene the housekeeping gene *GAPDH* was selected [136]. For control of individual host factor knockdowns primers were used employing the PrimerBank database [137, 138]. Primers for those as well as *GAPDH* are listed in Table S8. Primer concentration was 150 nM each, and primer efficiency was determined for each primer pair. RT-PCR was performed in an iCycler instrument (Bio-Rad, Munich, Germany) in triplicates in 96-well plates. Relative expression was determined using the 2^-ΔCt^ method [84, 139] with *GAPDH* expression set as 100%. Results were plotted using SigmaPlot 11 (Systat Software, Erkrath, Germany.

### Construction of plasmids

Plasmids used in this study were either obtained from Addgene, kind gifts from various laboratories, or cloned by Gibson Assembly or restriction enzyme digests and are listed in Table S6. Oligonucleotides for the construction of plasmids encoding host proteins fused to mRuby2 or EGFP are listed in Table S8. First, N- or C-terminal mRuby2 vectors were cloned. For that, the vectors pEGFP-C1 and pEGFP-N1 were amplified and EGFP was exchanged for a fragment encoding mRuby2. Genes encoding host proteins were amplified from vectors obtained from DNASU (Table S6) and then inserted into mRuby2 vectors by Gibson Assembly. Plasmids encoding host proteins fused to EGFP were constructed using restriction enzyme digests. The vector pEGFP-C3 was digested with *Kpn*I and *Xba*I or *Kpn*I and *Bam*HI and the larger fragment was recovered. The inserts were treated the same way and fragments were ligated.

### Host cell transfection

For LCI for localization of host factors, HeLa or HeLa-LAMP1-GFP cells were seeded 1 d prior to transfection. About 20.000 or 150.000 cells were seeded in 8-well chamber slides (see above) or 3.5 mm glass bottom dishes (FluoroDish, WPI, Berlin, Germany), respectively. For transfection 0.5 or 2 µg of plasmid DNA in 25 or 200 µL serum-free medium were mixed with 1 or 4 µL of FuGENE HD transfection reagent (DNA to reagent ratio of 1:2, Promega, Mannheim, Germany) and incubated for 10 min at RT. Medium on the cells was changed and transfection mixture applied. Cells were incubated for at least 18 h before infection with medium change during infection. For a complete list of transfection plasmids, see Table S6.

### Infection experiments

Overnight cultures of STM were diluted 1:31 and grown for additional 3.5 h in LB broth in glass test tubes with agitation in a roller drum. HeLa cells were infected with STM WT or *ssaV* for screening approaches in 96-well plates with a multiplicity of infection (MOI) of 15, otherwise for colocalization analysis or SIF quantification in 8-well chamber slides or FluoroDishes with an MOI of 75 or 50, respectively. Infection only of 96-well plates was synchronized by centrifugation at 500 x g for 5 min, and in all cases proceeded for 25 min at 37 °C in a humidified atmosphere containing 5% CO_2_. Cells were washed thrice with full medium or PBS for screening or non-screen LCI purposes, respectively, and incubated in full medium containing 100 µg/mL gentamicin for 1 h to eliminate extracellular bacteria. Then medium containing 10 µg/mL gentamicin was applied for the remainder of the experiment.

### Live cell imaging

For LCI full medium was replaced by imaging medium consisting of Minimal Essential Medium (MEM) with Earle’s salts, without NaHCO_3_, without L-glutamine and without phenol red (Biochrom, Berlin, Germany) supplemented with 30 mM HEPES (4-(2-hydroxyethyl)-1-piperazineethanesulfonic acid) (Sigma-Aldrich, Taufkirchen, Germany), pH 7.4, containing 10 µg/mL gentamicin. Fluorescence imaging for screening purposes was performed using a Zeiss Cell Observer microscope with Yokogawa Spinning Disk Unit CSU-X1 (Carl Zeiss, Göttingen, Germany), Evolve 512 x 512 EMCCD camera (Photometrics, Tucson, AZ, USA), automated PZ-2000 stage (Applied Scientific Instrumentation, Eugene, OR, USA), and infrared-based focus system Definite Focus, operated by Zeiss ZEN 2012 software (blue edition). The microscope was equipped with live cell periphery consisting of a custom-made incubation chamber surrounding the microscope body and connected with “The Cube” heating unit (Life Imaging Services, Basel, Switzerland) maintaining 37 °C and the Incubation System S for CO_2_ and humidity supply (PeCon, Erbach, Germany). Images were acquired using the Zeiss LD Plan-Neofluar 40x/0.6 Corr air objective (with bottom thickness correction ring). For acquisition of GFP and mCherry BP 525/50 (Zeiss) and LP 580 (Olympus, Hamburg, Germany) filters, respectively, were applied. All images obtained were processed by the ZEN software. Non-screen LCI was performed using a Leica SP5 confocal laser-scanning microscope (CLSM) operated by Leica LAS AF software. The microscope was also equipped with live cell periphery consisting of ‘The Box’ incubation chamber (Life Imaging Services, Basel, Switzerland), a custom-made heating unit and a gas supply unit ‘The Brick’. Images were acquired using the HCX PL APO CS 100x/1.4 oil objective (Leica, Wetzlar, Germany), applying the polychroic mirror TD 488/543/633 for acquisition of GFP and mCherry. All images were processed by LAS AF software.

### Quantification of SIF formation

After siRNA knockdown and infection, 100 infected HeLa-LAMP1-GFP cells per condition were examined live from 6-8 h p.i. for presence of SIF as exhibited by WT-infected cells, and percentage calculated. Results from biological triplicates were plotted using SigmaPlot 11.

### Data analysis

For central entry and collection of scoring data, the MATLAB-based utility SifScreen was used. Categorization of targets/hits was executed using the Gene Ontology classification scheme [140, 141]. For visualization of protein interactions, the STRING v10 database with default settings was applied [142].

## Suppl. Material

### Suppl. Tables

**Table S1.** Full list of the 496 host factors targeted in the siRNA screen including gene symbols, NCBI Gene IDs and accession numbers, full name, and aliases.

**Table S2.** Summary of the scoring of the executed siRNA screen with a full list of the trafficome, a list with hits only, and lists with low-, mid-, and high-ranking hits (scoring of 1-4, 5-7, or ≥8, respectively).

**Table S3.** Requirements of an RNAi screen with LCI for the identification of host factors involved in SIF formation by intracellular *Salmonella*.

| Type of prerequisite | Prerequisite | Necessity/Solution |
| --- | --- | --- |
| biological | • label for structure of interest<br>• *Salmonella* controls for structure of interest | • LAMP1 as marker of SIF<br>• WT: SIF-positive, *ssaV*: SIF-negative |
| technical | • fast and gentle microscopic system<br>• multiwell-compatible 37 °C incubation<br>• multiwell-compatible sufficient magnification<br>• autofocus system<br>• autofocus-compatible multiwell plates | • spinning disk confocal microscope (Zeiss)<br>• box incubation (custom-made)<br>• LD 40x/0.6 objective (+ bottom thickness correction collar, Zeiss)<br>• infrared-based system (Definite Focus, Zeiss)<br>• 96 Well Clear Bottom Black Cell Culture Microplates (0.5 mm bottom, Corning) |
| siRNA-related | • harmlessness of transfection reagent<br>• harmlessness of siRNA as reagent<br>• general efficacy of siRNA<br>• phenotype-interfering siRNA | • mock transfection control<br>• scrambled AllStars siRNA control<br>• lethal PLK1 siRNA control<br>• SIF-abolishing SKIP siRNA<br>• RT-PCR of SKIP knockdown |
|  | • control of phenotype-interfering siRNA |  |
| data analysis | • primary analysis | • manual visual inspection |
|  | • collection of analyzed data | • SifScreen utility |

**Table S4.**
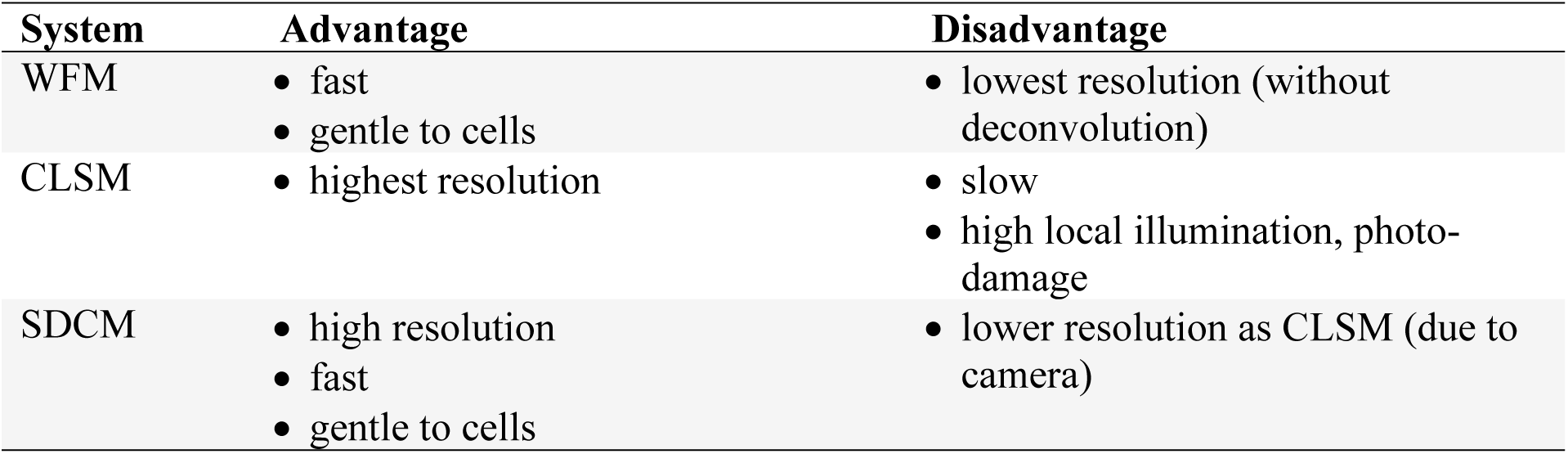
Characteristics of microscopy systems with respect to LCI and RNAi screening.

**Table S5.**
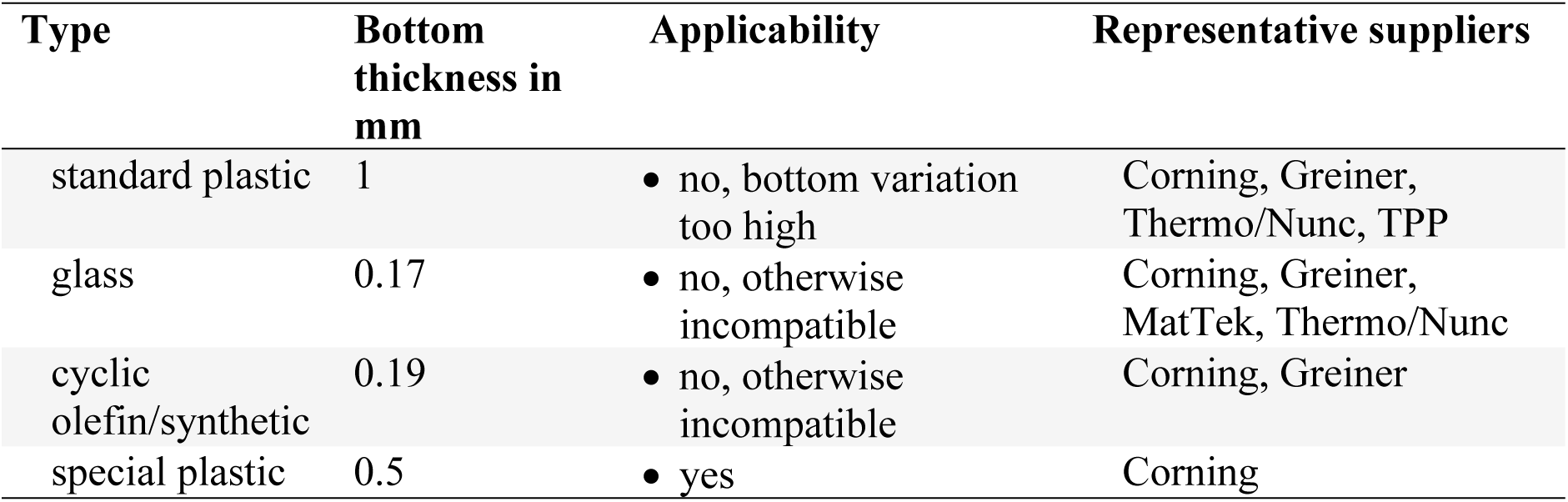
Suitability of various types of multi-well plates for screening with LD 40x objective and IRF.

**Table S6.**
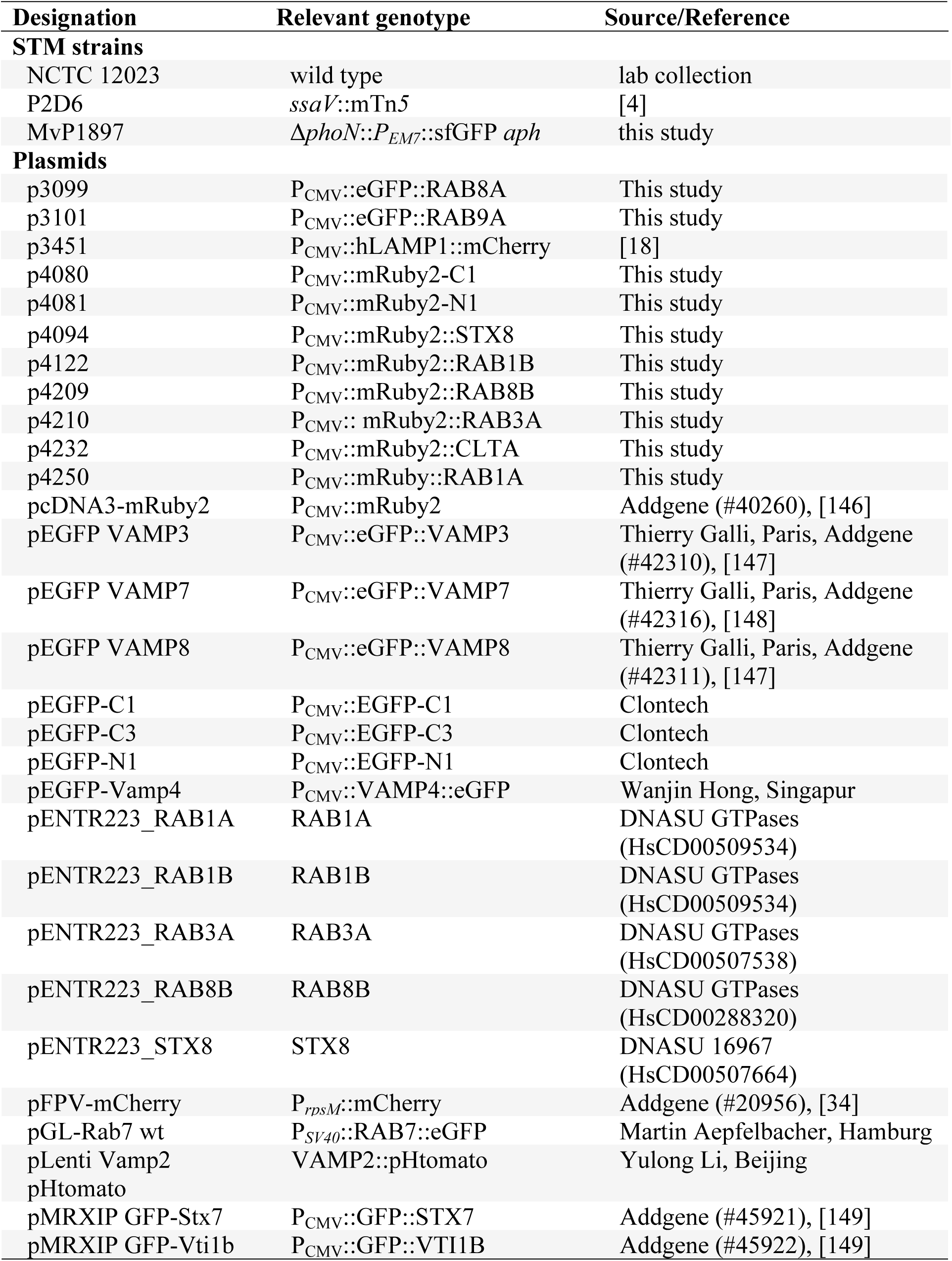
Bacterial strains and plasmids used in this study.

**Table S7.**
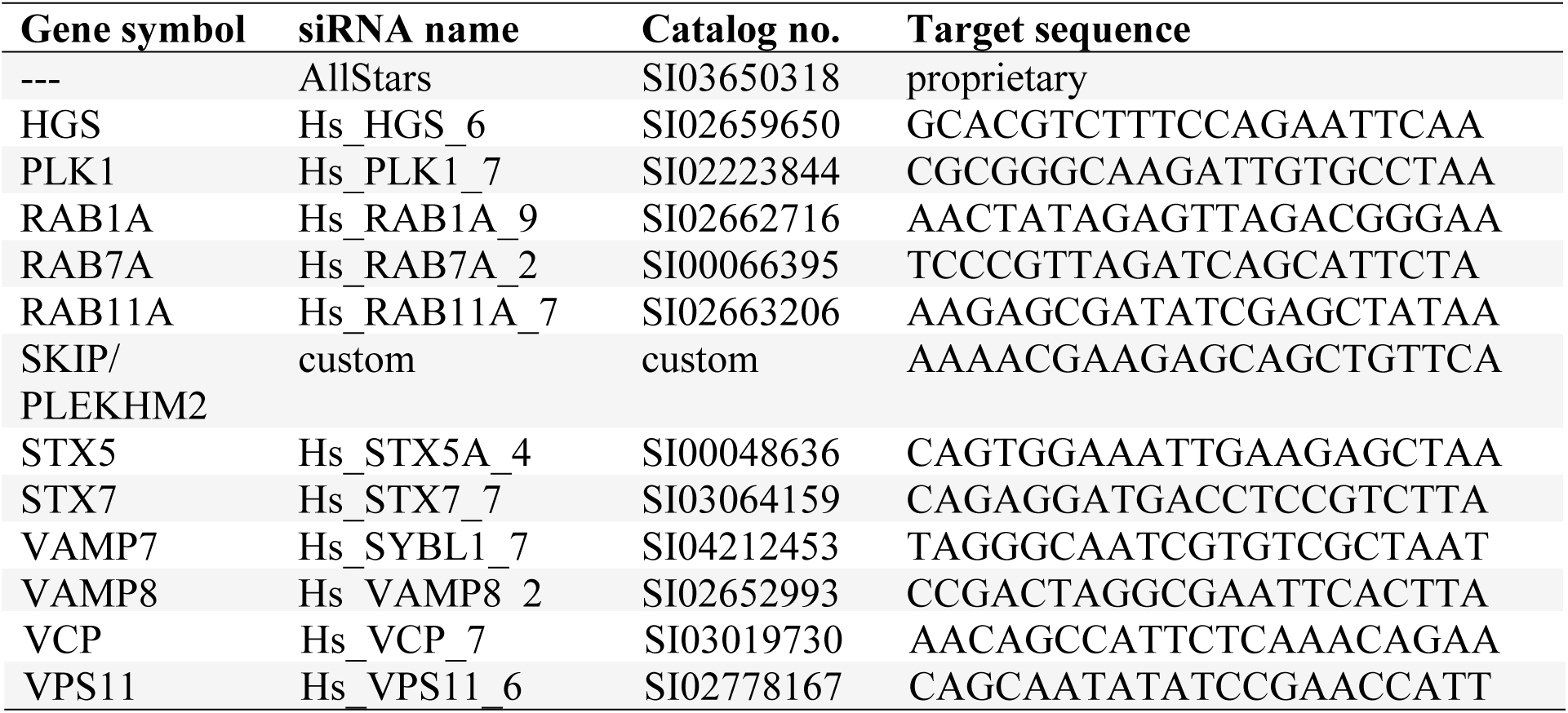
Individual siRNA information used for validation.

**Table S8.**
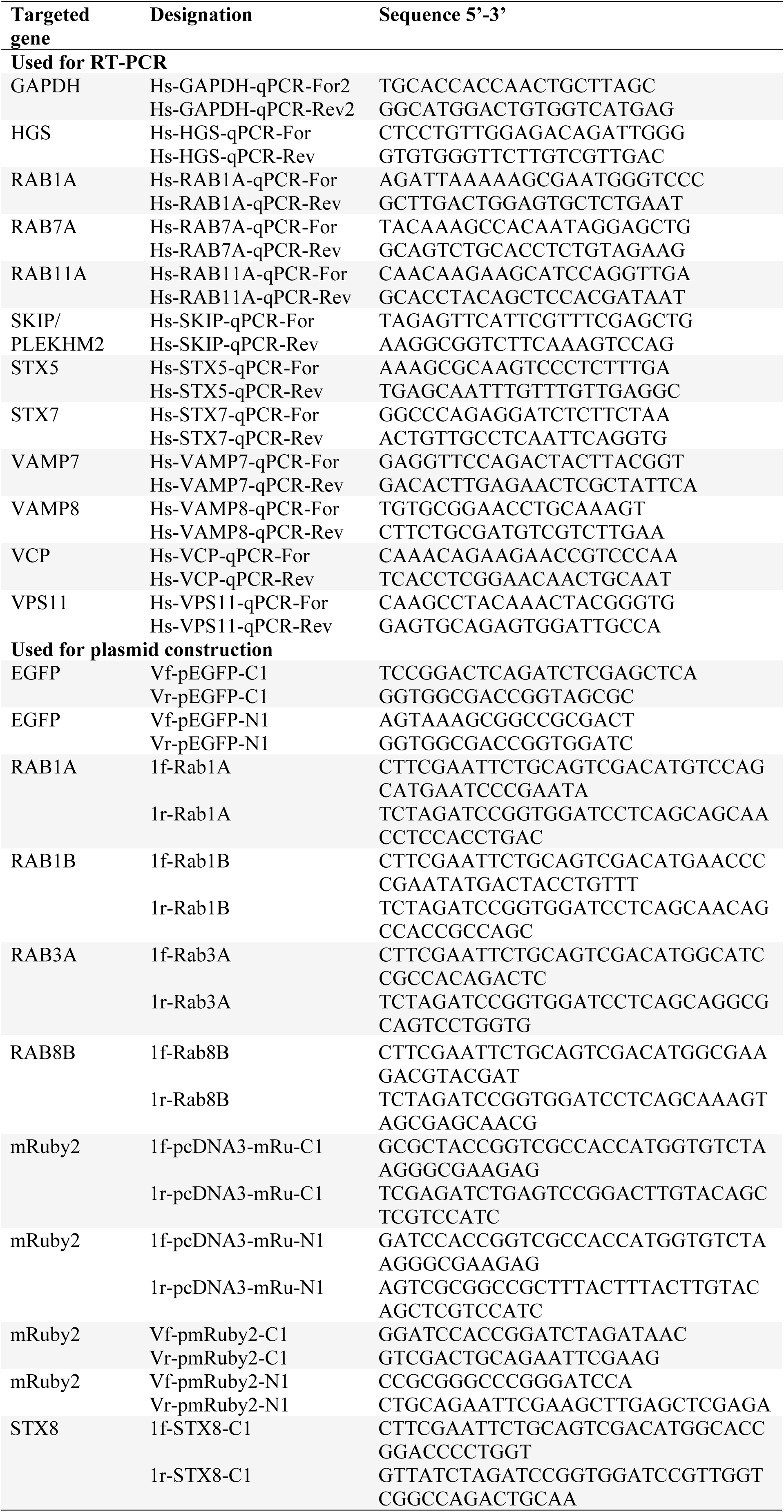
Oligonucleotides in this study.

### Suppl. Figure legends

**Figure S1. Validation of host factor siRNA silencing.** HeLa-LAMP1-GFP cells were reverse transfected with siAllStars or the indicated siRNA. Total RNA was extracted, mRNA reverse transcribed, and the generated cDNA was used in RT-PCR. Depicted are means with standard deviation for three biological replicates (*n* = 3) each performed in triplicates. Statistical analysis was performed against siAllstars with Student’s *t*-test and indicated as: ***, *p* < 0.001.

### Suppl. Movie captions

**Movie 1. Time-lapse acquisition of infected siAllStars-treated cells.** The movie corresponds to Figure 1D.

**Movie 2. Time-lapse acquisition of infected siSKIP-treated cells.** The movie corresponds to Figure 1D.

